# Improved Resolution of Highly Pathogenic Avian Influenza Virus Haemagglutinin Cleavage Site Using Oxford Nanopore R10 Sequencing Chemistry

**DOI:** 10.1101/2023.09.30.560331

**Authors:** Jeremy D Ratcliff, Brian Merritt, Hannah Gooden, Jurre Y Siegers, Abhi Srikanth, Sokhoun Yann, Sonita Kol, Sarath Sin, Songha Tok, Erik A Karlsson, Peter M Thielen

## Abstract

Highly pathogenic avian influenza viruses continue to pose global risks to One Health, including agriculture, public, and animal health. Rapid and accurate genomic surveillance is critical for monitoring viral mutations, tracing transmission, and guiding interventions in near real-time. Oxford Nanopore sequencing holds promise for real-time influenza genotyping, but data quality from R9 chemistry has limited its adoption due to challenges resolving low-complexity regions such as the biologically critical hemagglutinin cleavage site, a homopolymer of basic amino acids that distinguish highly pathogenic strains. In this study, human and avian influenza isolates (n=45) from Cambodia were sequenced using both R9.4.1 and R10.4.1 flow cells and chemistries to evaluate performance between approaches. Overall, R10.4.1 yielded increased data output with higher average quality compared to R9.4.1, producing improved consensus sequences using a reference-based bioinformatics approach. R10.4.1 had significantly lower minor population insertion and deletion frequencies, driven by improved performance in low sequence complexity regions prone to insertion and deletion errors, such as homopolymers. Within the hemagglutinin cleavage site, R10.4.1 resolved the correct motif in 90% of genomes compared to only 60% with R9.4.1. Further examination showed reduced frameshift mutations in consensus sequences generated with R10.4.1 that could result in incorrectly classified virulence. Improved consensus genome quality from nanopore sequencing approaches, especially across biologically important low-complexity regions, is critical to reduce subjective hand-curation and will improve local and global genomic surveillance responses.

## Introduction

As the coronavirus disease 2019 (COVID-19) pandemic response wanes, expanded global sequencing capacity is now primed to analyze other pathogens of concern. Rapid whole genome sequencing enables real-time molecular epidemiology to track viral evolution, trace transmission pathways, guide prevention and intervention strategies during seasonal circulation and outbreaks, and develop new medical countermeasures ^1–6^. Recent advancements in third-generation sequencing technologies, such as Oxford Nanopore Technologies (ONT) platforms, have promised a revolution in the field of viral genomics by enabling real-time, long-read sequencing with portable and cost-effective devices ^7^. This next generation of sequencing has facilitated rapid genomic characterization critical for response to viruses during outbreaks, including those caused by Ebola, Zika, and severe acute respiratory syndrome coronavirus 2 (SARS-CoV-2), as well as endemic viruses such as Dengue and Chikungunya ^8–12^. Oxford Nanopore long-read sequencing shows promise for field epidemiology of priority pathogens, but concerns over platform accuracy have limited its adoption for routine influenza genomic surveillance ^8^.

Genomic surveillance is required to address the One Health risk posed by Influenza A viruses, which threaten global public health, animal health, and food security. Avian influenza viruses (AIV) cause severe poultry outbreaks and remain an important zoonotic threat ^13^. Since 2020, highly pathogenic (HP) AIV, mainly A/H5N1 clade 2.3.4.4b, has caused an unprecedented number of deaths in wild birds and poultry and, continues to spread into mammalian species, including marine life and domestic animals. ^14,15^ Occasional spillover and human cases continue annually, including recent fatal HPAIV A/H5Nx infections as well as low pathogenicity (LP) AIV A/H9N2 in Cambodia and China ^16–18^. ONT sequencing utility has been demonstrated for whole genome sequencing from cultured isolates and clinical specimens. ^19–24^ In addition to human seasonal surveillance, these techniques have been important in influenza surveillance and response - including in field deployable versions ^25,26^ - in avian species ^27,28^, swine ^26,29^, and AIV spillover into humans ^17^. However, the hemagglutinin (HA) cleavage site of avian influenza, a major virulence determinant, is composed of basic amino acids encoded by a low-complexity A/G-rich portion of the genome. Resolution of these low-complexity regions (LCRs) has been a challenge for many sequencing platforms, and has limited the adoption of nanopore sequencing for widespread HPAIV genomic surveillance ^20^. Accurately sequencing LCRs is important for many pathogens relevant to human and animal health ^30–33^.

Beginning in late 2023, Oxford Nanopore will transition to a new sequencing chemistry (R10) that is being described as having improved overall data quality over R9 chemistry, which was initially released in 2016. As this new chemistry has been introduced, independent validations of performance have shown variable results ^34,35^. In HPAIV, the hemagglutinin (HA) protein contains a multibasic cleavage site upstream of an RGLF amino acid motif with a homologous sequence of adjacent arginine (R) and lysine (K) residues. This region acts as a key virulence factor that allows avian influenza viruses to replicate to a high titer systemically in birds ^36^. Given the critical importance and biological significance of this highly homopolymeric multibasic cleavage site in influenza ^37^, and the increased likelihood of sequencing issues near homopolymer regions ^38^, it is essential to understand how R10 chemistry can improve ONT sequencing for rapid, comprehensive, accurate, and cost-effective genomic surveillance of influenza viruses, especially AIV. This study provides a timely assessment of Oxford Nanopore’s new R10.4.1 sequencing chemistry compared to the widely used R9.4.1 for avian influenza characterization. Comparative analysis of influenza viruses (n=46), with a focus on HPAIV, was conducted to evaluate the performance improvement between chemistries. Improved influenza genome resolution, especially across the problematic hemagglutinin cleavage site homopolymer, enables more reliable and rapid genomic surveillance to inform outbreak response. By evaluating sequencing accuracy gains on biological samples, this work aims to address a key evidence gap that will help guide the adoption of ONT sequencing for molecular epidemiology of priority zoonotic diseases by increasing overall confidence in data produced by the platform.

## Methods

### Ethics Approval

Animal sampling was conducted in conjunction with the National Animal Health and Production Research Institute (NAHPRI) under the direction of the General Directorate for Animal Health and Production, Cambodian Ministry of Agriculture, Forestry and Fisheries (MAFF) as part of routine disease surveillance activities; thus, poultry sampling was not considered experimental animal research. The analysis of poultry samples for avian influenza testing was approved by the Cambodian National Ethics Committee for Health Research (approval 2022/#044NECHR). The Institut Pasteur du Cambodge (IPC) serves as the National Influenza Center in Cambodia and World Health Organization H5 Regional Reference Laboratory, with approvals and infrastructure necessary to work on highly pathogenic avian influenza. No animal experimentation was performed at IPC.

### Sample Collection and Processing

Active AIV surveillance in Cambodia was performed by Institut Pasteur du Cambodge (IPC) in collaboration with the NAHPRI under the direction of the General Directorate for Animal Health and Production, MAFF. Briefly, oropharyngeal and cloacal samples were collected from domestic birds (ducks and chickens) as described previously ^39^. Human samples are collected regularly through the influenza-like illness (ILI) and severe acute respiratory illness (SARI) surveillance systems in Cambodia and sent to IPC for isolation and characterization ^40,41^. Viral RNA was extracted using the QIAamp Viral RNA Mini Kit (Qiagen, Maryland, USA) according to the manufacturer’s protocol. All samples were screened by real-time reverse transcription PCR targeting the M gene, as previously described ^39,41^. Samples with a cycle threshold (Ct) less than 40 were deemed positive. Avian-derived samples with Ct<30 were used for isolation under biological safety level 3 conditions using 10-day-old embryonated chicken eggs inoculated via the allantoic route and were harvested 72 hrs later. Positive human samples were isolated on Madin-Darby Canine Kidney (MDCK) cells under enhanced biological safety level 2 conditions as previously described ^42^. The presence of influenza virus in the supernatant and allantoic fluid was tested using a hemagglutination (HA) assay ^43^. Both original samples and isolates were subtyped using influenza RT-PCR assays to test for human seasonal and avian subtypes ^28,41,44^. In total, 45 unique isolates were sequenced for this study, with seasonal human influenza used as positive controls (Table S1). Isolates were randomly selected from active, longitudinal surveillance with a focus on available low-pathogenic and highly pathogenic strains.

### Sanger Sequencing

To enable Sanger sequencing, the HA gene of A/H5Nx viruses was amplified using conventional PCR as described previously ^45^. Briefly, HA were amplified using cycling conditions: 5 min at 94°C followed by 35 cycles of amplification with denaturation at 95°C for 30 sec. annealing at 50°C for 30 sec, extension at 72°C for 2 min and a final extension of 10 min at 72°C. Primer pairs used were H5F-1: AGCAAAAGCAGGGGTYTAAT, H5R-1111rmod: CCATACCAACCATCTAYCATTC, H5F-800: TTATWGCTCCYGAATATGCATACAA, and H5R-1710: AGTCAAATTCTGCATTGTAACGACC. PCR product was visualized on 1.5% agarose gel by electrophoresis. PCR products were sent to Macrogen (https://dna.macrogen.com/), South Korea for Sanger sequencing.

### Nanopore Sequencing

Whole genomes were amplified using a modified multi-segment RT-PCR approach that incorporates integrated molecular indices, with a degenerate base at position four of the Uni-12 primer (5’AGC**<u>R</u>**AAAGCAGG) ^28,46^. Resulting indexed RT-PCR products were pooled and prepared for sequencing using ligation sequencing kit SQK-LSK109 (R9 chemistry) or SQK-LSK114 (R10 chemistry), and sequenced on the GridION platform (Oxford Nanopore Technologies, Oxford, UK). Final library concentrations were 35.4ng/μl and 28.2ng/μl for SQK-LSK109 and SQK-LSK114 reagent kits, respectively. Molarity was determined using Oxford Nanopore recommended tables, arbitrarily assuming an average multi-segment RT-PCR fragment size of 2kb. For 20 femtomole (fmol) libraries, 26 nanograms (26 ng = 20 fmol) of the final DNA library were loaded onto flowcells. For the 140 fmol libraries, 180 nanograms (180 ng = 138 fmol) of the final DNA library were loaded onto flowcells.

### Bioinformatic processing

Sanger data was assembled using Geneious® 9.0.5 (https://www.geneious.com) to generate a consensus sequence. ONT data were basecalled and demultiplexed using Oxford Nanopore Guppy v6.4.6 (ONT), with high accuracy basecalling enabled and custom indexing configured for integrated indexing MS-PCR. Demultiplexed reads were assembled using the CDC Iterative Refinement Meta-Assembler (IRMA) version 1.0.3 module for Flu-Minion ^47^. A total of 42 gene segment sequences obtained in this study were deposited at the GISAID database under accession number EPI2666609-EPI2666644. Unprocessed sequence data were deposited to the NCBI Sequence Read Archive under bioproject SUB13867526.

### Nanopore performance assessment

Various metrics were used to assess the performance of R9 and R10 chemistries. For calculation of sequence quality, only reads that mapped to influenza segments (as identified from the IRMA output) were considered. These values were derived from the sequencing summary file. Read length distribution was taken from the raw FASTQ file. Breadth of X coverage and sequencing depth at individual timepoints were calculated using pysam v0.21.0 ^48^ as mapped to each segment in the reference output of the IRMA v1.03 pipeline. Individual read sequencing times were pulled from the sequencing summary file generated after Guppy basecalling. Time to 95% coverage was calculated by sorting each of threads on the start time from the sequencing summary’s metadata and identifying when the corresponding value of bases, identified in the pileup subcommand of pysam, has been reported.

### Local Sequence Complexity

For a given genome position, consensus-level local sequence complexity was defined as the mean Shannon entropy value for the five component 5mers that include the position of measure (from -4 to +4). Shannon entropy values for 5mers were calculated using the formula: 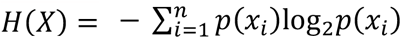, where n is the total number of unique bases in the 5mer input and p(xi) is the proportion of all bases of a given type in the 5mer. Perfect homopolymers (e.g., “AAAAA”) return a Shannon entropy value of 0. Values were derived in python3 leveraging Biopython, Pandas, and user-defined functions ^49,50^.

### Per-position and 9mer indel rate calculations

Per-position indel rates were defined as the total number of mapped indels per-position in the IRMA-defined consensus sequence divided by the total coverage for that position. BAM files for each genome segment were analyzed using Pysamstats ^51^. Per-position coverage was defined as the sum of mapped matches, mismatches, insertions, and deletions to an index. Analyses and aggregations were limited to positions that had a coverage depth >100X based on the Pysamstats output, which calculates statistics against genome positions (i.e., each row in the output is a position in the consensus sequence).

After limiting by coverage depth, total counts of matches, mismatches, insertions, and deletions were aggregated for each unique 9mer in the consensus sequences of an individual sample. Briefly, for each Pysamstats output the first and final four positions of each segment were discarded (as they are not at the center of 9mers). Then, the Pysamstats outputs for each sample were appended into a single data file and all unique 9mers observed for that sample were recorded. For each unique 9mer in the merged Pysamstats output, sample-level indel rates were calculated by summing aggregate coverage and indel counts for each segment. This approach results in per-sample rate estimates weighted by total coverage within a sample.

### Evaluation of nanopore performance on individual 9mers

Global indel rate estimates for individual 9mers were calculated as the mean of per-sample estimates within the respective chemistry. These were not corrected for variations in coverage between samples. Only 9mers observed in three or more samples were included in analyses. When comparing R9 and R10 chemistries, the standard error of the difference of means was calculated using the formula: 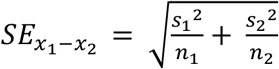. These standard error values were used to derive confidence interval (CI) estimates for the difference of means. To ensure robustness given the large number of 9mers under investigation, significant differences in performance between R9 and R10 were defined as those 9mers whose difference of means 99% CI did not cross 0.

### Identification of multibasic cleavage site

The diversity of indels throughout the HA segment prevented discrimination of the multibasic cleavage solely through nucleotide indices. To identify the multibasic cleavage site, the consensus sequence for the HA segment from the sanger output for the 21 H5NX samples was translated using Pysam and Biopython ^49,50^. A regular expression ((R|K)[^RKH](R|K)) was used to search the amino acid sequence for the location of a multibasic cleavage site. After identification, consensus sequence outputs for R9 and R10 chemistries were aligned to the Sanger sequencing references manually. Alignments were visually inspected for substitutions, insertions, and deletions.

## Results

### R10 chemistry outperforms R9 on multiple metrics

This study characterized the comparative performance of leveraging Oxford Nanopore R10 sequencing chemistry in place of R9 chemistry for sequencing influenza A virus samples and, in a subset of HPAIV samples, Sanger sequencing of the HA multibasic cleavage site (Figure 1). The 46 samples sequenced include both human and avian influenza viruses of multiple subtypes from circulating human seasonal and avian influenza viruses collected in Cambodia (Table S1).

**Figure 1:**
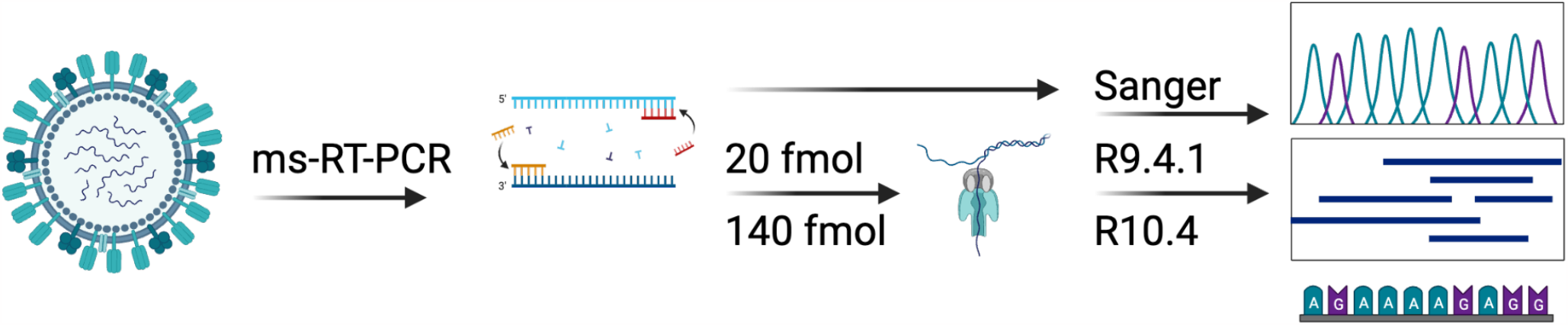
Experimental Validation of Sequencing Chemistry Performance for Avian Influenza Viruses. 46 Influenza A virus samples were subjected to multi-segment (ms) PCR and sequenced on either a Sanger or Oxford Nanopore platform. For samples sequenced using Oxford Nanopore chemistry, both R9 and R10 sequencing chemistries were evaluated. The resulting nanopore data were then analyzed using the IRMA software analysis pipeline to evaluate the impact of sequencing chemistry on a consensus generation pipeline.

Recommended DNA loading concentrations for Oxford Nanopore influenza sequencing range between 20-50fmol ^52^. To evaluate the impact of increased concentrations on data output and quality, flowcells were loaded at 20 and 140fmol on both R9 and R10 flowcells. A substantial increase in overall data output was observed when loading flowcells with 140fmol, with no apparent reduction in sequencing output over the 24-hours during which sequencing was performed (Figure 2A). Across all timepoints (1 minute intervals), 140fmol loading concentrations increased the output (measured in Gbp) for R10 chemistry by an average of 48% and R9 chemistry by an average of 575% versus a 20fmol input. For the R9 chemistry, this increase in data output also resulted in a higher proportion of genome segments being recovered across all samples (Figure 2B). Complete genome segments were recovered more frequently, regardless of sequence length, when sequencing with R10 flow cells loaded at 140fmol compared to either R9 loading concentration. However, limited improvements were observed after ten hours of run time for all but PB1 and PB2 segments (Figure S1).

**Figure 2:**
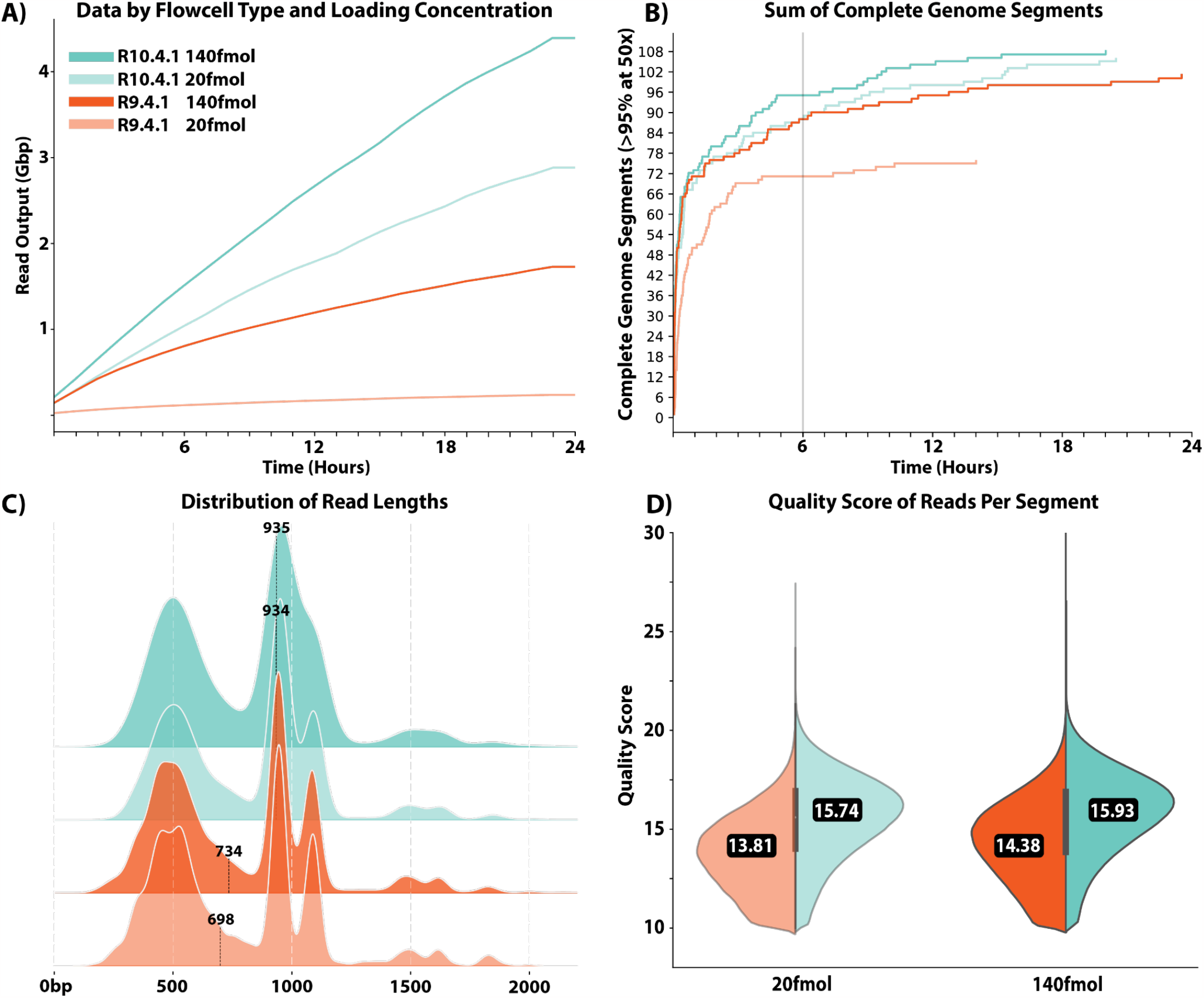
Performance metrics of R9 and R10 sequencing chemistries. Figures display data generated from 46 samples. (A) Total read output in Gbp over time for R9 and R10 sequencing chemistries and both loading concentrations. (B) Total number of complete gene segments across all samples that have reached >95% coverage at a sequencing depth of at least 50X at a given time. All gene segments are treated equally. C and D) Distribution of read lengths and quality scores, respectively, across all reads that mapped to the consensus IRMA output. Values displayed in each plot indicate medians.

Ultimately, the trends observed across sequencing chemistries and loading concentrations indicate a higher relative coverage rate for all eight segments for both the low- and high-concentration loading conditions for R10 flowcells. While varying degrees of coverage relative to segment type were observed, the overall findings show that R10 is the preferred chemistry at nearly all time points of the sequencing runs performed, with the highest disparity for R10 between segments NS, MP, and NA (Figure S1). While the prevalence of the depth of genome coverage varies across both runs and segments, we see a general consistency in the speed of achieved thresholds for R10 compared to R9.

Median read quality scores and read lengths were evaluated after remapping individual reads to the segment reference output from IRMA. Median read quality scores were significantly higher for R10 than R9 (p < 5e-324 for 20fmol and 140fmol, two-tailed Mann-Whitney U tests; Figure 2D). Using R10 in place of R9 resulted in an average increase in read accuracy of 1.35% for 20fmol (97.21% vs 95.86%) and 1.02% for 140fmol (97.33% vs 96.31%). Increasing the loading concentrations also resulted in slight, but significant, improvements in average quality for both chemistries (p < 5e-324 in both instances). Of note, a minimal proportion of reads from either R9 (0.13% for 20fmol and 0.32% for 140fmol) or R10 (0.83% for 20fmol and 1.07% for 140fmol) achieved Q20 quality. Despite libraries being generated from the same amplicon pool, the distribution of read lengths was significantly different between R10 and R9 chemistries. R10 consistently produced significantly longer reads (p < 5e-324 for 20fmol and 140fmol, two-tailed Mann-Whitney U tests; Figure 2C). R10 reads were, on average, 33.8% longer than R9 at the 20fmol loading concentration and 27.4% longer at the 140fmol loading concentration. Increased loading concentration resulted in significantly longer reads for R9 (p = 4.35e-56) and for R10 (p = 3.21e-47), albeit the magnitude of impact on read length was minimal for R10 (medians of 934 vs 935 base pairs).

### Increased deletion rate is associated with decreased local sequence complexity in both R9 and R10 flow cells

Nanopore-based sequencing has been criticized for high rates of sequencing error in homopolymeric regions that are characterized by minimal sequence complexity ^53^. These sequencing errors often present as insertions or deletions in individual reads relative to the consensus sequence. To investigate the role of local sequence complexity in observed indel rates, per-position complexity values were compared to minor population insertion and deletion rates observed in the IRMA output. Per-position complexity was defined as the average of the Shannon entropy values for the five 5mers that contain the position of interest in the consensus sequence. 5mers were chosen as this is the oligonucleotide length that occupies the pore in R7 chemistries and determines the signal in R9 chemistry, although the number of bases that influence the ONT raw signal is somewhat variable and dependent on the nanopore chemistry used ^54,55^. Through this method, all reads mapped to a specific position are inferred to be derived from the cumulative signal across all five composite 5mers. For the following sections analyzing 9mer-level indel rates, only the 140fmol samples were considered for statistical independence.

There are 262,144 possible combinations for an oligonucleotide of nine bases. Across all samples in this study, a total of 42,654 unique 9mers were observed, of which 29,705 were brought forward for analysis after filtering for detection in three or more samples. Unsurprisingly, the observed 9mers had a slightly but significantly lower distribution of Shannon entropy values than the unobserved 9mers (medians of 1.46 vs 1.49, p = 4.7e-257, two-tailed Mann-Whitney U Test). Further, the included 9mers had a significantly lower distribution of Shannon entropy values than those filtered out (n = 12,949), albeit the median values for both populations w<u>ere</u> 1.46 (p = 4.0e-5, two-tailed Mann-Whitney U test) (Figure S2).

For both R9 and R10 flow cells, 9mer sequence complexity was significantly negatively correlated with average deletion rate (p =1.3e-251, linear regression, Figure S2). Less complex segments of the genome had higher observed deletion rates. The parameter estimate was 5.6x larger in R9 than R10 (-0.023 vs -0.004). Curiously, insertion rates were significantly negatively correlated with sequence complexity in R9 (-0.001, p =5.0e-13) and significantly positively correlated in R10 (0.001, p = 1.7e-11, Figure S2), although the magnitude of the effect was minimal for both chemistries.

It was separately observed that per-position total coverage was significantly negatively correlated with 9mer insertion and deletion rates for both chemistries (p <0.002 in all instances, linear regression). This finding is intuitive, as a single errant insertion or deletion read in a low coverage position would have a larger effect on observed indel rates than in a high coverage position. Furthermore, mean complexity values were positively correlated with coverage (p = 0.0002 for both R9 and R10, linear regression), suggesting that the observed associations between indel rates and sequence complexity may simply be a mediator of coverage. However, the significant relationships and directions of effect between sequence complexity and indel rates remained for both chemistries when controlled for coverage (p < 4.5e-10 in all instances, linear regression).

### Global improvement in deletion rate in R10 chemistry is driven by improved performance in LCRs

To characterize the driver of the improved indel rates in R10 chemistry, we evaluated the change in indel rate when changing chemistries for each 9mer using difference of means analysis. Of the 29,705 included 9mers, 8,441 (28.4%) had significantly lower deletion rates in R10 while only 1,050 (3.5%) had significantly lower deletion rates in R9 (Figure 3A, D). Among those with significant differences, the magnitude of improvement was significantly different between the two chemistries, with those favoring R10 improving by a median of 1.2% and those favoring R9 improving by a median of 1.0% (p = 3.82e-9, two-tailed Mann-Whitney U Test; Figure 3B, E).

**Figure 3:**
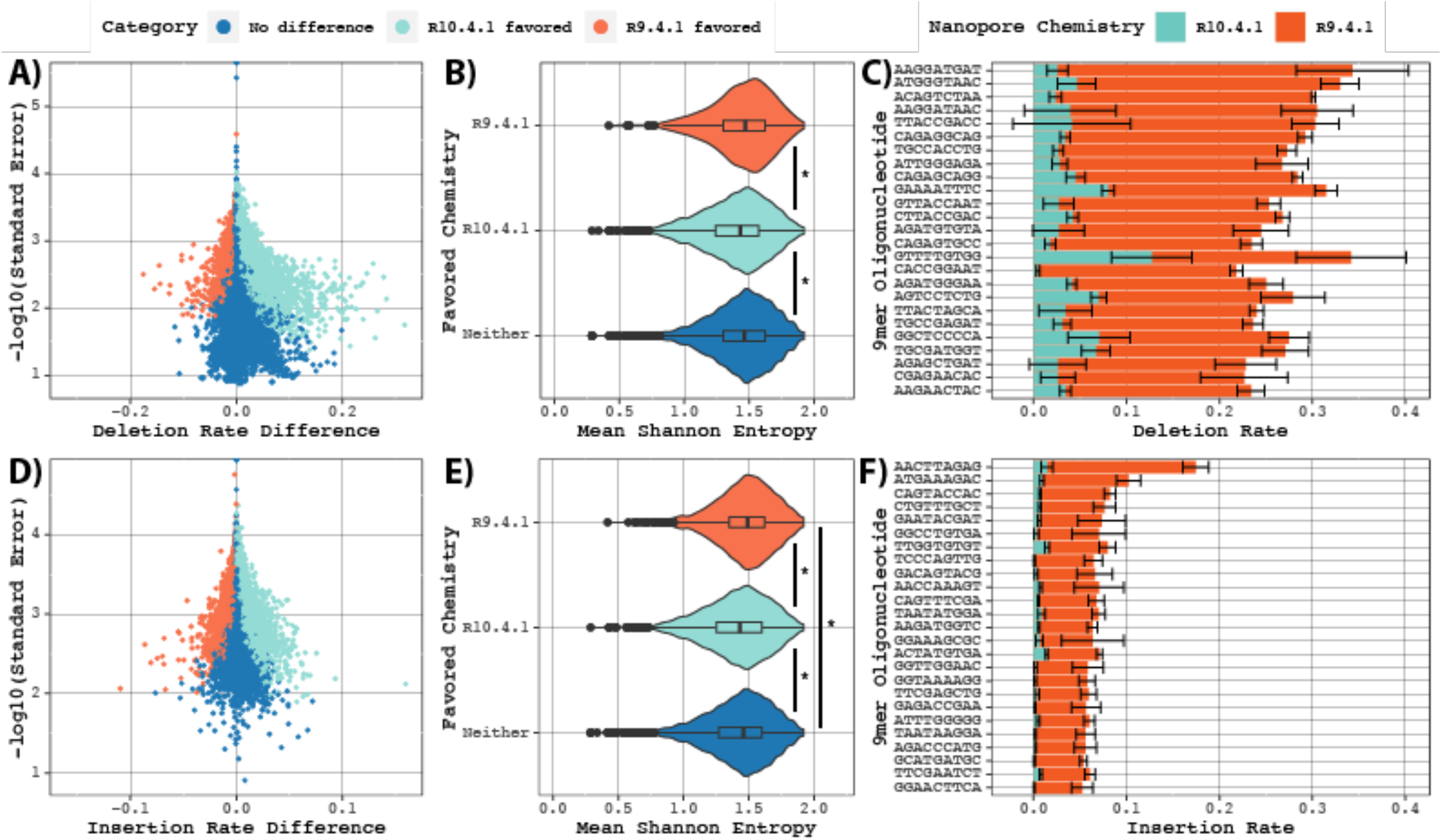
Reduced Insertion and Deletion Rates in R10 Chemistry. Improvements in using R10 chemistry in place of R9 for individual 9mers were observed in three or more samples. Favored chemistries determined by significant differences in difference of means analysis. A,D) Volcano plots of individual 9mers, colored based on favored chemistry. B/E) Violin plots of mean Shannon entropy values for 9mers within each favored chemistry. Boxplots display median and first and third quartiles. Asterisks display population differences significant by two-tailed Mann-Whitney U Test. C,F) 9mers with the top 25 significant reduction in indel rates moving from R9 to R10 chemistry for deletions and insertions, respectively. Error bars display the mean ± one standard deviation.

Improvements in insertion rates were less clearly weighted towards R10, with 8,591 (28.9%) and 2,932 (9.9%) having significant improvements in R10 and R9, respectively. There was no significant difference in magnitude of improvement in insertion rate between the two chemistries (medians of 0.42% and 0.43%, respectively, p = 0.843, two-tailed Mann-Whitney U Test).

From the observation that sequence complexity had a smaller influence on observed deletion rate in R10 than R9, it was hypothesized that sequence complexity may partially explain the favored chemistry for individual 9mers. In regards to deletion rates, 9mers that favored R10 had significantly lower mean Shannon entropy values than those that favored R9 or neither (medians of 1.43, 1.47, and 1.46, respectively, p values of 9.48e-9 and 2.43e-34, two-tailed Mann-Whitney U Tests). There was no difference between those without a favored chemistry and those that favored R9 (p = 0.36, two-tailed Mann-Whitney U Test). Further, mean Shannon entropy values were a significant predictor of the difference between R10 and R9 estimates among all 9mers (p < 2e-308, linear regression). These results strongly suggest that the overall reduction in deletion rate in R10 can be ascribed to increased performance in low-complexity 9mers.

The relationship was similar for insertions, although in this instance there were significant differences between all three groups (p < 1.5e-6 in all instances, two-tailed Mann-Whitney U Tests; Figure 3 C, F). Median Shannon entropy values were lowest in R10 (1.43), followed by those without a favored chemistry (1.46), and those that favored R9 (1.49). Similarly, mean Shannon entropy values predicted the difference in insertion rates (p = 9.3e-47). Other factors, such as base composition (see supplementary results), may also partially explain the categorization of individual 9mers. Overall, these results demonstrate that the global reduction in indel rates in R10 chemistry relative to R9 can largely be ascribed to improved performance in low-complexity 9mers.

### R10 chemistry Characterization of homopolymer region at the hemagglutinin cleavage site

To characterize the performance of R9 and R10 sequencing chemistry to accurately discriminate the multibasic cleavage site, an LCR, the consensus level outputs for the HA segment were aligned and the adenine homopolymer upstream of the RGLF motif was scrutinized. Consistent with the higher observed indel rate in R9 vs R10 chemistry, both R9 20fmol and 140fmol performed worse than both loading concentrations of R10 at accurately resolving the length of this homopolymer (Figure 4B). At the recommended loading concentration of 20fmol, R10 was significantly more likely than R9 to generate the correct consensus length for the homopolymer (p = 0.0076, two-sample proportion test). This significant improvement remained when aggregating data for both loading concentrations, with R9 correctly identifying the 5mer in only 59.5% of samples compared to 90.4% for R10 (p = 0.0025, two-sample proportion test). For the R9 chemistry, all of the detected indels resulted in a frameshift mutation. For R10, only one of the two detected indels disrupted the RGLF motif, with one three-base insertion only lengthening the multibasic motif. Upon realignment of data to a Sanger reference, these insertions appear to result from strand-specific errors in R9 data, and from improper reference identification errors in both data types (Figure S3).

**Figure 4:**
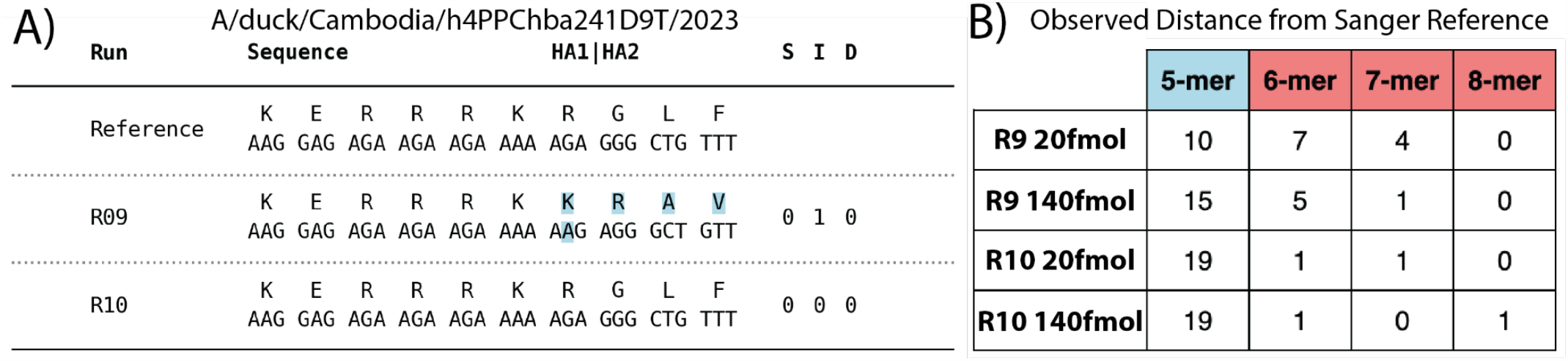
Improved Resolution of the H5 Multi-Basic Cleavage Site. (A) Representative multiple sequence alignment for HA segment inclusive of the multibasic cleavage motif, composed of Sanger (Reference), R9, and R10 consensus outputs. 20fmol and 140fmol consensus outputs were identical for both R9 and R10 chemistries for this sample. R9 output includes a single base insertion at the 3’ end of the adenine homopolymer upstream of the RGLF motif, resulting in an apparent frameshift mutation (highlighted in blue). S = # of substitutions / I = # of insertions / D = # of deletions. (B) Length of adenine homopolymer in consensus outputs for both chemistries and loading concentrations for all high pathogenicity avian influenza samples (n = 21). Lengths greater than five bases are assumed to be artifactual insertions.

## Discussion

The question is not if, but when, the next influenza pandemic will occur. Rapid genomic surveillance is critical for monitoring avian influenza virus evolution and mitigating pandemic risks. Long-read nanopore sequencing enables real-time whole genome characterization to inform outbreak response, but data accuracy concerns have slowed adoption. This study demonstrates that Oxford Nanopore’s new R10 chemistry significantly improves sequencing performance for avian influenza viruses compared to the widely used R9, particularly in problematic homopolymeric regions like the virulence-determining hemagglutinin cleavage site.

Across all segments, R10 chemistry yielded faster sequencing output, higher coverage uniformity, longer reads, and reduced error rates versus R9. The lower insertion and deletion frequencies with R10 were driven by improved resolution of repetitive motifs prone to sequencing errors, like homopolymers. At the hemagglutinin cleavage site, a homopolymer of basic amino acids distinguishing high pathogenicity strains, R10 resolved the correct motif in 90% of samples compared to just 60% with R9. Further examination revealed fewer R10-induced frameshifts that could misclassify virulence in automated analysis pipelines. By increasing accuracy in avian influenza genomes, especially across critical LCRs, the R10 nanopore chemistry facilitates timely and reliable genomic surveillance to inform outbreak response.

The enhanced accuracy of influenza sequencing demonstrated here could substantially reduce costs and delays hindering large-scale genomic epidemiology of avian influenza and other priority zoonoses. Generating reliable genomes faster directly decreases labor and supplies needed ^8^, as well as the turnaround time from sampling to actionable data ^10^. With improved data quality, less effort is required for troubleshooting or confirmation testing, further cutting costs. Operationally, faster and more accurate on-site sequencing with Oxford Nanopore systems reduces reliance on centralized sequencing facilities, lowering transport expenses and turnaround times ^26^. By scaling up capacity without sacrificing reliability, routine genomic surveillance becomes more feasible globally ^23^. While quantification is difficult, even incremental technical gains can produce outsized impacts on overcoming financial and logistical barriers to genomic epidemiology programs ^8^.

Beyond economic benefits, faster and higher confidence avian influenza sequencing data provides key health protections. Accuracy is critical for surveillance systems to detect emerging infectious diseases and distinguish them from known pathogens ^56^. Each day of delayed pandemic response could increase eventual cases by 3%, so earlier outbreak detection and characterization is critical ^57^. At the start of the COVID-19 pandemic, many groups were hesitant to submit data due to questionable accuracy, especially given the low mutation rate of SARS-CoV-2 when there were few mutations observed as the pandemic became more widespread ^58^. As confidence in bioinformatics methods increased, substantially more data became available for global genomic surveillance activities due to improved informatics capabilities ^59^. Delayed reporting of the initial H5N1 human cases in Vietnam in 2003-2004 similarly delayed public health interventions and contributed to limited containment ^60^. Enhanced genomic situational awareness would also better inform interventions like culling high-risk poultry flocks or closing markets, while tracing transmission pathways more quickly to limit spread. Closing live poultry markets within a week of case detection during the 2013-2014 H7N9 outbreak in China reduced human infections by 99% compared to closures greater than a week post-detection ^61^. Overall, transitioning to R10 nanopore chemistry will directly strengthen avian influenza surveillance and response capabilities for both agriculture and public health.

This study has several limitations related to the samples and methods utilized, and we anticipate this comparative dataset will be useful for exploring additional benefits and shortcomings of nanopore sequencing chemistries. The integrated indexing multi-segment PCR approach has not been thoroughly evaluated alongside standard ms-RT-PCR methods, and may introduce unintended segment amplification biases ^62^. Evaluation of ms-RT-PCR product size distributions additionally presents challenges for evaluating molar concentration of a sequencing library, and read lengths observed in sequencing data did not match initial assumptions during flowcell loading. In addition, only a subset of laboratory-cultured A/H5Nx avian influenza viruses were analyzed for hemagglutinin cleavage site resolution accuracy, so other subtypes and direct clinical, poultry, or environmental samples may require alternative analysis to characterize the improvements for specific motifs. Further, the limited dataset and consensus-based bioinformatics analysis leave many additional sequence features unaddressed, such as minor variants or *de novo* assembly challenges.

The approach used to calculate indel rates based on minor variants will misassign indels that impact the consensus sequence (i.e., ‘proper’ reads on an artifactual consensus-level deletion will appear as insertions upon minor variant analysis). Bioinformatics pipelines for reference-based influenza consensus generation were restricted to the “gold standard” for influenza sequencing, IRMA, and it is possible that future versions of this software or alternative computational approaches, such as Oxford Nanopore’s Medaka, would produce improved influenza reference sequences if properly integrated into the analysis pipeline ^47,63^. While substantial gains were demonstrated for avian influenza sequencing when using R10 sequencing chemistry, further assessment across sample types, basecalling models, and additional pathogens is warranted to confirm the expanded benefits of transitioning to these new methodologies.

## Conclusions

The threat of the next influenza pandemic underscores the need for timely and accurate genomic surveillance. While Oxford Nanopore sequencing has shown utility for real-time influenza monitoring, issues with data quality have limited widespread adoption. This study demonstrates that the new R10 chemistry significantly improves sequencing accuracy compared to R9, particularly in problematic homopolymeric regions like the multibasic cleavage site of highly pathogenic avian influenza viruses. Though sample preparation and bioinformatic pipelines still potentially need further refinement to reduce issues, R10 chemistry enables faster production of higher confidence influenza genomes from field and clinical samples. By increasing data reliability, the R10 chemistry facilitates timely situational awareness and informed response during outbreaks. With continuous improvements to nanopore sequencing, Oxford Nanopore platforms are approaching the speed, portability, and accuracy needed for routine influenza genomic epidemiology in humans and animals globally.

## References

1. Ghedin, E. et al. Large-scale sequencing of human influenza reveals the dynamic nature of viral genome evolution. Nature 437, 1162–1166 (2005).

2. Rambaut, A. et al. The genomic and epidemiological dynamics of human influenza A virus. Nature 453, 615–619 (2008).

3. Wille, M. & Barr, I. G. Resurgence of avian influenza virus. Science 376, 459–460 (2022).

4. Hoque, M. A., Burgess, G. W., Cheam, A. L. & Skerratt, L. F. Epidemiology of avian influenza in wild aquatic birds in a biosecurity hotspot, North Queensland, Australia. Prev. Vet. Med. 118, 169–181 (2015).

5. Liu, J. et al. Highly pathogenic H5N1 influenza virus infection in migratory birds. Science 309, 1206 (2005).

6. Rafique, S. et al. Global review of the H5N8 avian influenza virus subtype. Front. Microbiol. 14, 1200681 (2023).

7. Gardy, J. L. & Loman, N. J. Towards a genomics-informed, real-time, global pathogen surveillance system. Nat. Rev. Genet. 19, 9–20 (2018).

8. Kafetzopoulou, L. E. et al. Assessment of metagenomic Nanopore and Illumina sequencing for recovering whole genome sequences of chikungunya and dengue viruses directly from clinical samples. Euro Surveill. 23, (2018).

9. Fauver, J. R. et al. Coast-to-Coast Spread of SARS-CoV-2 during the Early Epidemic in the United States. Cell 181, 990–996.e5 (2020).

10. Quick, J. et al. Real-time, portable genome sequencing for Ebola surveillance. Nature 530, 228–232 (2016).

11. Quick, J. et al. Multiplex PCR method for MinION and Illumina sequencing of Zika and other virus genomes directly from clinical samples. Nat. Protoc. 12, 1261–1276 (2017).

12. Faria, N. R. et al. Establishment and cryptic transmission of Zika virus in Brazil and the Americas. Nature 546, 406–410 (2017).

13. Ongoing avian influenza outbreaks in animals pose risk to humans. https://www.who.int/news/item/12-07-2023-ongoing-avian-influenza-outbreaks-in-animals-pose-risk-to-humans.

14. Leguia, M. et al. Highly pathogenic avian influenza A (H5N1) in marine mammals and seabirds in Peru. Nat. Commun. 14, 5489 (2023).

15. Rabalski, L. et al. Emergence and potential transmission route of avian influenza A (H5N1) virus in domestic cats in Poland, June 2023. Euro Surveill. 28, (2023).

16. Surveillance Division of the Communicable Disease Branch of the Centre for Health Protection. Avian Influenza Report. Reporting period: Aug 20, 2023 – Aug 26, 2023 (Week 34). Center for Health Protection - Avian Influenza Report https://www.chp.gov.hk/files/pdf/2023_avian_influenza_report_vol19_wk34.pdf (2023).

17. Um, S. et al. Human Infection with Avian Influenza A(H9N2) Virus, Cambodia, February 2021. Emerg. Infect. Dis. 27, 2742–2745 (2021).

18. World Health Orgainzation. Situation Report: Human infection with avian influenza A(H5) viruses (18 August 2023). WHO Surveillance Report - Avian Influenza https://www.who.int/westernpacific/emergencies/surveillance/avian-influenza (2023).

19. Wang, J., Moore, N. E., Deng, Y.-M., Eccles, D. A. & Hall, R. J. MinION nanopore sequencing of an influenza genome. Front. Microbiol. 6, 766 (2015).

20. Yip, C. C.-Y. et al. Nanopore Sequencing Reveals Novel Targets for Detection and Surveillance of Human and Avian Influenza A Viruses. J. Clin. Microbiol. 58, (2020).

21. Keller, M. W. et al. Direct RNA Sequencing of the Coding Complete Influenza A Virus Genome. Sci. Rep. 8, 14408 (2018).

22. Crossley, B. M. et al. Nanopore sequencing as a rapid tool for identification and pathotyping of avian influenza A viruses. J. Vet. Diagn. Invest. 33, 253–260 (2021).

23. Ip, H. S., Uhm, S., Killian, M. L. & Torchetti, M. K. An Evaluation of Avian Influenza Virus Whole-Genome Sequencing Approaches Using Nanopore Technology. Microorganisms 11, (2023).

24. King, J., Harder, T., Beer, M. & Pohlmann, A. Rapid multiplex MinION nanopore sequencing workflow for Influenza A viruses. BMC Infect. Dis. 20, 648 (2020).

25. de Vries, E. M. et al. Rapid, in-field deployable, avian influenza virus haemagglutinin characterisation tool using MinION technology. Sci. Rep. 12, 11886 (2022).

26. Rambo-Martin, B. L. et al. Influenza A Virus Field Surveillance at a Swine-Human Interface. mSphere 5, (2020).

27. Miah, M. et al. Culture-Independent Workflow for Nanopore MinION-Based Sequencing of Influenza A Virus. Microbiol Spectr 11, e0494622 (2023).

28. Edwards, K. M. et al. Detection of Clade 2.3.4.4b Avian Influenza A(H5N8) Virus in Cambodia, 2021. Emerg. Infect. Dis. 29, 170–174 (2023).

29. Vereecke, N. et al. Successful Whole Genome Nanopore Sequencing of Swine Influenza A Virus (swIAV) Directly from Oral Fluids Collected in Polish Pig Herds. Viruses 15, (2023).

30. Mier, P. & Andrade-Navarro, M. A. The Conservation of Low Complexity Regions in Bacterial Proteins Depends on the Pathogenicity of the Strain and Subcellular Location of the Protein. Genes 12, (2021).

31. María Velasco, A. et al. Low complexity regions (LCRs) contribute to the hypervariability of the HIV-1 gp120 protein. J. Theor. Biol. 338, 80–86 (2013).

32. Chaudhry, S. R., Lwin, N., Phelan, D., Escalante, A. A. & Battistuzzi, F. U. Comparative analysis of low complexity regions in Plasmodia. Sci. Rep. 8, 335 (2018).

33. Monzon, S., Varona, S., Negredo, A. & Patiño-Galindo, J. A. Changes in a new type of genomic accordion may open the pallets to increased monkeypox transmissibility. bioRxiv (2022).

34. González-Recio, O. et al. Sequencing of SARS-CoV-2 genome using different nanopore chemistries. Appl. Microbiol. Biotechnol. 105, 3225–3234 (2021).

35. Luo, J. et al. Systematic benchmarking of nanopore Q20+ kit in SARS-CoV-2 whole genome sequencing. Front. Microbiol. 13, 973367 (2022).

36. Cox, M. M. & Nelson, D. L. Lehninger Principles of Biochemistry. vol. 5 (unknown, 2000).

37. Kawaoka, Y. & Webster, R. G. Sequence requirements for cleavage activation of influenza virus hemagglutinin expressed in mammalian cells. Proc. Natl. Acad. Sci. U. S. A. 85, 324–328 (1988).

38. Van den Hoecke, S., Verhelst, J., Vuylsteke, M. & Saelens, X. Analysis of the genetic diversity of influenza A viruses using next-generation DNA sequencing. BMC Genomics 16, 79 (2015).

39. Suttie, A. et al. Diversity of A(H5N1) clade 2.3.2.1c avian influenza viruses with evidence of reassortment in Cambodia, 2014-2016. PLoS One 14, e0226108 (2019).

40. Horwood, P. F. et al. Circulation and characterization of seasonal influenza viruses in Cambodia, 2012-2015. Influenza Other Respi. Viruses 13, 465–476 (2019).

41. Siegers, J. Y. et al. Genetic and Antigenic Characterization of an Influenza A(H3N2) Outbreak in Cambodia and the Greater Mekong Subregion during the COVID-19 Pandemic, 2020. J. Virol. 95, e0126721 (2021).

42. Horm, S. V. et al. Epidemiological and virological characteristics of influenza viruses circulating in Cambodia from 2009 to 2011. PLoS One 9, e110713 (2014).

43. Killian, M. L. Hemagglutination assay for influenza virus. Methods Mol. Biol. 1161, 3–9 (2014).

44. Horwood, P. F. et al. Co-circulation of Influenza A H5, H7, and H9 Viruses and Co-infected Poultry in Live Bird Markets, Cambodia. Emerg. Infect. Dis. 24, 352–355 (2018).

45. Al-Qahtani, A. A. et al. Characterization of H5N1 influenza A virus that caused the first highly pathogenic avian influenza outbreak in Saudi Arabia. J. Infect. Dev. Ctries. 9, 1210–1219 (2015).

46. Thielen, P. Influenza Whole Genome Sequencing with Integrated Indexing on Oxford Nanopore Platforms. (2022).

47. Shepard, S. S. et al. Viral deep sequencing needs an adaptive approach: IRMA, the iterative refinement meta-assembler. BMC Genomics 17, 708 (2016).

48. pysam: Pysam is a Python module for reading and manipulating SAM/BAM/VCF/BCF files. It’s a lightweight wrapper of the htslib C-API, the same one that powers samtools, bcftools, and tabix. (Github).

49. Cock, P. J. A. et al. Biopython: freely available Python tools for computational molecular biology and bioinformatics. Bioinformatics 25, 1422–1423 (2009).

50. The pandas development team. pandas-dev/pandas: Pandas. (2023). doi:10.5281/zenodo.7857418.

51. Miles, A. pysamstats: A fast Python and command-line utility for extracting simple statistics against genome positions based on sequence alignments from a SAM or BAM file. (Github).

52. Ligation sequencing influenza whole genome V14. Ligation sequencing influenza whole genome V14 (VERSION: INF_9189_V114_REVB_14JUN2023) https://community.nanoporetech.com/docs/prepare/library_prep_protocols/ligation-sequencing-influenza-whole-genome-v14/v/inf_9189_v114_revc_14jun2023 (2023).

53. Delahaye, C. & Nicolas, J. Sequencing DNA with nanopores: Troubles and biases. PLoS One 16, e0257521 (2021).

54. Rang, F. J., Kloosterman, W. P. & de Ridder, J. From squiggle to basepair: computational approaches for improving nanopore sequencing read accuracy. Genome Biol. 19, 90 (2018).

55. Wang, Y., Zhao, Y., Bollas, A., Wang, Y. & Au, K. F. Nanopore sequencing technology, bioinformatics and applications. Nat. Biotechnol. 39, 1348–1365 (2021).

56. Iuliano, A. D. et al. Estimates of global seasonal influenza-associated respiratory mortality: a modelling study. Lancet 391, 1285–1300 (2018).

57. Ferguson, N. M. et al. Strategies for containing an emerging influenza pandemic in Southeast Asia. Nature 437, 209–214 (2005).

58. Hodcroft, E. B. et al. Want to track pandemic variants faster? Fix the bioinformatics bottleneck. Nature Publishing Group UK http://dx.doi.org/10.1038/d41586-021-00525-x (2021) doi:10.1038/d41586-021-00525-x.

59. Nicholls, S. M. et al. CLIMB-COVID: continuous integration supporting decentralised sequencing for SARS-CoV-2 genomic surveillance. Genome Biol. 22, 196 (2021).

60. Normile, D. Avian influenza. Vietnam battles bird flu … and critics. Science 309, 368–373 (2005).

61. Chen, Y., Cheng, J., Xu, Z., Hu, W. & Lu, J. Live poultry market closure and avian influenza A (H7N9) infection in cities of China, 2013-2017: an ecological study. BMC Infect. Dis. 20, 369 (2020).

62. Zhou, B. et al. Single-reaction genomic amplification accelerates sequencing and vaccine production for classical and Swine origin human influenza a viruses. J. Virol. 83, 10309–10313 (2009).

63. Lee, J. Y. et al. Comparative evaluation of Nanopore polishing tools for microbial genome assembly and polishing strategies for downstream analysis. Sci. Rep. 11, 20740 (2021).

